# Complementary studies of lipid membrane dynamics using iSCAT and super-resolved Fluorescence Correlation Spectroscopy

**DOI:** 10.1101/235564

**Authors:** Francesco Reina, Silvia Galiani, Dilip Shrestha, Erdinc Sezgin, Gabrielle de Wit, Daniel Cole, B. Christoffer Lagerholm, Philip Kukura, Christian Eggeling

**Affiliations:** MRC Human Immunology Unit and Wolfson Imaging Centre Oxford, Weatherall Institute of Molecular Medicine, University of Oxford, Oxford, United Kingdom; Physical and Theoretical Chemistry Laboratory, Department of Chemistry, University of Oxford, Oxford, United Kingdom; Wolfson Imaging Centre Oxford, Weatherall Institute of Molecular Medicine, University of Oxford, Oxford, United Kingdom; Institute of Applied Optics, Friedrich-Schiller University and Leibniz Institute of Photonic Technology (IPHT), Jena, Germany; Leibniz Institute of Photonic Technology (IPHT), Jena, Germany

## Abstract

Observation techniques with high spatial and temporal resolution, such as single-particle tracking (SPT) based on interferometric Scattering (iSCAT) microscopy, and fluorescence correlation spectroscopy applied on a super-resolution STED microscope (STED-FCS), have revealed new insights of the molecular organization of membranes. While delivering complementary information, there are still distinct differences between these techniques, most prominently the use of fluorescent dye-tagged probes for STED-FCS and a need for larger scattering gold nanoparticle tags for iSCAT. In this work we have used lipid analogues tagged with a hybrid fluorescent tag – gold nanoparticle construct, to directly compare the results from STED-FCS and iSCAT measurements of phospholipid diffusion on a homogeneous Supported Lipid Bilayer (SLB). These comparative measurements showed that while the mode of diffusion remained free, at least at the spatial (>40 nm) and temporal (50 ≤ t ≤ 100 ms) scales probed, the diffussion coefficient was reduced by 20- to 60-fold when tagging with 20 and 40 nm large gold particles as compared to when using dye-tagged lipid analogues. These FCS measurements of hybrid fluorescent tag – gold nanoparticle labeled lipids also revealed that commercially supplied streptavidin-coated gold nanoparticles contain large quantities of free streptavidin. Finally, the values of apparent diffusion coefficients obtained by STED-FCS and iSCAT differed by a factor of 2-3 across the techniques, while relative differences in mobility between different species of lipid analogues considered were identical in both approaches. In conclusion, our experiments reveal that large and potentially crosslinking scattering tags introduce a significant slow-down in diffusion on SLBs but no additional bias, and our labeling approach creates a new way of exploiting complementary information from STED-FCS and iSCAT measurements.

## Introduction

The study of cellular membrane dynamics is a subject of intense research in Biophysics, due to its relevance for many cellular processes, such as cellular signaling. Important questions in this field revolve around the functionality of membrane-associated protein-protein and protein-lipid interactions, for example, the formation and the function of lipid nanodomains or “rafts” [1]. These structures, due to their sub-diffraction limit size (< 200 nm) and transient nature (< milliseconds), have so far proven challenging to identify and study *in vivo* [2]. Therefore, biophysical studies often focus on observing molecular diffusion dynamics on model membranes such as Supported Lipid Bilayers (SLBs) or synthetic and cell-derived vesicles, aiming to transfer or compare results to the *in vivo* case (e.g. [3]–[5]). Two prominent methods for observing molecular diffusion on membranes are Fluorescence Correlation Spectroscopy (FCS) [6], [7] and single-particle tracking (SPT) [8], [9].

In FCS, fluctuations of the fluorescence intensity from a single observation spot are recorded. These are mainly due to the movement of fluorescent molecules diffusing in the sample. The autocorrelation function of such fluctuations is analyzed through a suitable model to determine the characteristics of the motion of the diffusing molecules. In particular, it is possible to derive parameters like the average transit time through the observation spot and (if the dimension of the spot is known) also the apparent diffusion coefficient D. However, FCS experiments recorded on conventional far-field microscopes limit the minimal size of the observation spot, due to the diffraction limit of light, and this inevitably leads to that nanoscopic (< 150-200 nm) obstacles or heterogeneities in diffusion are missed [7]. It is possible to overcome this limitation either by extrapolating the results from diffraction-limited FCS recordings to the nanoscale [10], or by combining super-resolution STED microscopy with FCS (STED-FCS) [11],[12]. The latter provides a direct method for tuning the observation spot size of FCS [12], [13], and therefore allows direct monitoring of molecular dynamics at sub-diffraction length scales (30-200nm). This has revealed the presence of new complex anomalous diffusion modes at small spatial and short time scales [11]. Nevertheless, a complete understanding of diffusion-related phenomena requires complementary observation approaches. In effect, while (STED-)FCS provides robust statistics on dynamical properties of specific molecular populations, it might miss certain aspects of diffusion heterogeneity due to the inherent averaging in the calculation of the auto-correlation function. Therefore, molecular populations with different diffusion characteristics, or molecules changing their diffusion mode over time could remain undetected. Accessing this level of detail requires direct tracking of molecular motions over space and time, as in Single Particle Tracking (SPT) [8], [9]. In a typical SPT experiment, a series of large field-of-view images of individual isolated molecules are acquired and their spatial positions localized and followed over time. While SPT experiments have highlighted important features of molecular motions, they are ultimatively limited by issues such as low signal-to-noise ratios, low temporal resolution and photobleaching, which in turn leads to limited recording times and short trajectories.

Recently developed Interferometric Scattering (iSCAT) microscopy-SPT has shown exceptional localization precision and temporal resolution [14], [15] and is thus a good candidate to complement STED-FCS. Image detection in iSCAT microscopy relies on the interference of coherent light scattered by objects in the sample with the reflected component of the same beam. Using this approach, it is possible to detect seconds-long trajectories of single particles as they move through the sample, with sampling rates of up to 500kHz and localization precision in the nanometer or sub-nanometer range [16]. This has made it possible to disclose, with so far unprecedented detail, the inhomogeneity of lipid diffusion in model membranes, and also transient confinement events, which cannot be revealed purely by relying on conventional SPT methods such as Mean Squared Displacement [17]–[19]. A major limitation of SPT techniques such as iSCAT, however, is that they often require large (10-40nm in diameter) [20] scattering tags, such as gold nanoparticles. The diagram in Figure 1a illustrates this issue by comparing the relative dimensions of streptavidin-coated, 40 nm-diameter gold nanoparticles (often used in iSCAT and related SPT approaches [9]) and of a fluorescent dye tagged lipid analogue (as used in STED-FCS). The gold nanoparticles are on one hand several orders of magnitude larger than the target lipid and on the other hand coated with a variable number of multivalent streptavidin molecules. A combination of their large size and the potential to crosslink many target molecules may hinder the motion of the target molecule [21]. On one hand, size-dependent hydrodynamics affect the diffusion of the whole lipid-gold nanoparticle system since the nanoparticle, which is positioned near a “wall” constituted by the lipid bilayer, has a diffusion coefficient that is determined by a modified Stokes-Einstein equation, several orders of magnitude slower than a lipid in a Supported Lipid Bilayer [22]. On the other hand, the large gold nanoparticles usually express several binding sites and may therefor crosslink several lipids, an effect that is however not easily quantifiable, given the stochastic nature of the phenomenon, and the numerous factors that can influence it, like the density of the binding-site coating. In contrast, the size of a fluorescent dye is comparable to that of the diffusing lipid. Consequently, a side-by-side comparison of both approaches, i.e. iSCAT-based SPT against STED-FCS and gold-nanoparticle vs fluorescence dye tagging, would highlight possible artefacts in either method. Such comparison has been performed before for other techniques such as SPT and image-correlation spectroscopy methods [23], FCS and FRAP (Fluorescence Recovery after Photobleaching) [24], or FRAP and SPT [25]. The comparison between SPT and STED-FCS has thus far only been limited to discuss the principles of the analysis approaches [26], without considering direct experimental comparison for data from the same sample (e.g. gold nanoparticle and fluorescent dye tagged lipid reporters). This is however relevant since iSCAT- (or generally gold nanoparticle-) based SPT and STED-FCS experiments may reach the same spatial and temporal scales. The main reason for the lack of the latter experiments lies in that the respective lipid analogues could only be employed in either approach, i.e. dye-tagged lipid analogues in STED-FCS and gold nanoparticle tagged lipid analogues in iSCAT. While the dye-tagged analogues do not produce sufficient scattering contrast compared to unlabelled lipids in iSCAT (necessitating gold beads), the gold nanoparticle tagged lipids were not useful for STED-FCS due to missing fluorescence signal (necessitating fluorescent dyes).

**Figure 1.**
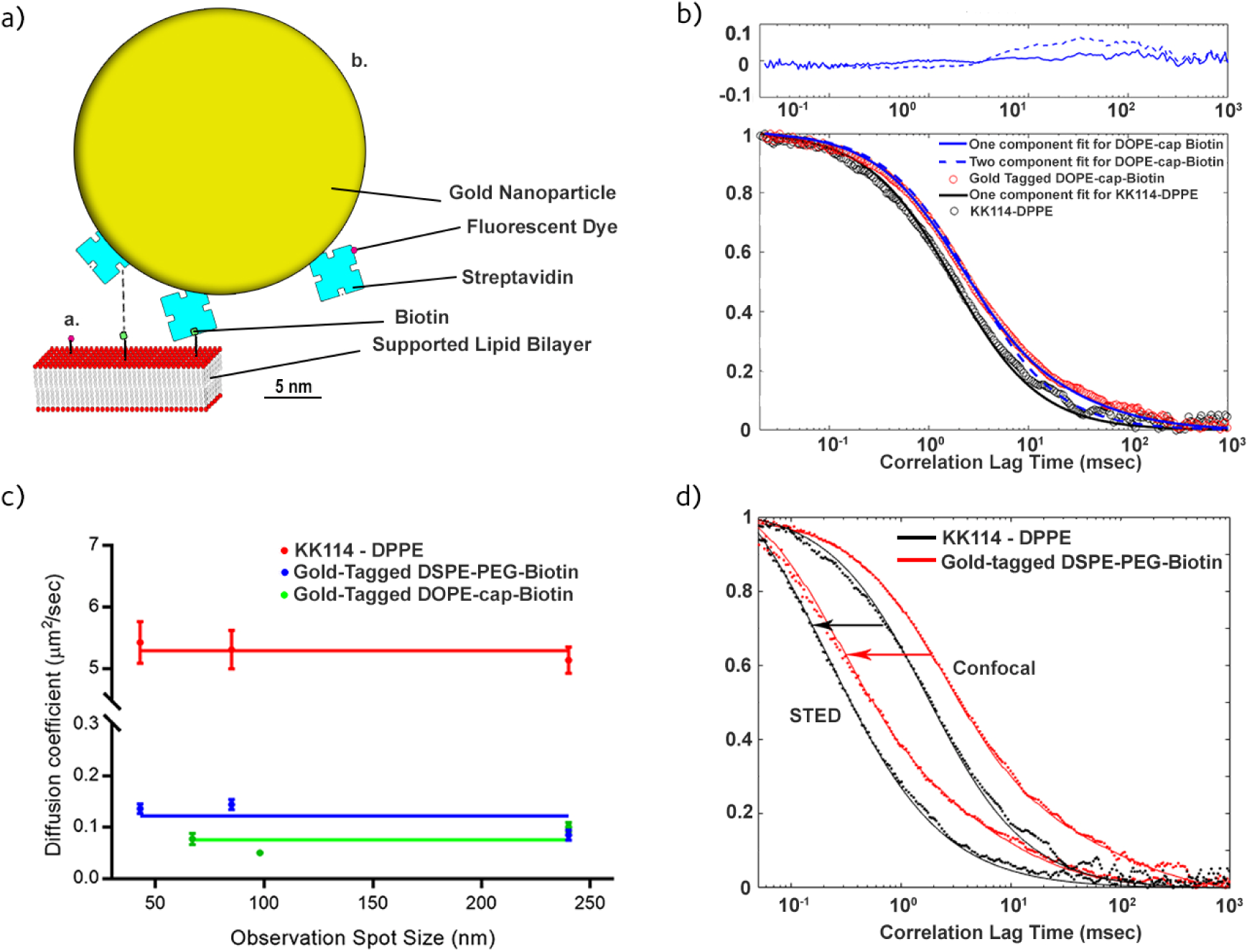
a) Schematic scale presentation of the employed fluorescent lipid analogues in the lipid bilayer (red: lipid head groups, grey: lipid chains): Tagged with i) small organic dye (red), and ii) with a 40nm gold nanoparticle (yellow) coated with organic-dye (red) tagged streptavidin (blue) binding to a biotinylated (green) lipid, possibly introducing cross-linking to a second biotinylated lipid (dashed line). b) Representative confocal FCS data taken on the homogeneous DOPC SLB for the fluorescent gold nanoparticle-tagged DSPE-PEG-Biotin lipid (red circles) with one- (blue dashed line) and two-component fits (solid blue line), and the fluorescent lipid analogue KK114-DPPE (black circles) with one-component fit (black line). Upper panel: Residuals of the one-(dashed line) and two-component (solid line) fits to the FCS data taken for the fluorescent gold nanoparticle tagged lipid analogue. c) Dependence of the apparent diffusion coefficient D on the observation spot diameter d as taken from the STED-FCS recordings of the different fluorescent analogues, KK114-tagged DPPE (red), gold nanoparticle tagged DSPE-PEG-Biotin (blue) and gold nanoparticle tagged DOPE-cap-Biotin (green) with the average value plotted across the D values as guide for the eyes (solid lines). Mean values (dots) and error bars (standard deviation) from n≥8 measurements. d) Representative confocal and STED-FCS data (as labelled, STED for d = 85nm) taken for the KK114-tagged DPPE (black dots) and fluorescent gold nanoparticle tagged DSPE-PEG-Biotin (red dots) in the SLBs, and two-component fits to the data, respectively (solid lines).

In this work, we present a protocol for attaching a fluorescent dye to streptavidin-coated gold nanoparticles, thus making them usable both as fluorescent and scattering tags. Using STED-FCS, the diffusion characteristics of such gold nanoparticle tagged phospholipids were compared to that of organic-dye tagged phospholipid analogues. While the diffusion mode of the gold-tagged phospholipids remained unaltered (i.e., free, as expected of phospholipids on a fluid SLB) by the mere presence of the tag, their mobility was reduced by a factor of roughly 20 to 60 (depending on the particle size) compared to the dye-only tagged lipids. This phenomenon is most likely due to tag-induced cross-linking of several lipids and, probably less effectively, by the size of the nanoparticles themselves. Yet, their diffusion remained free as expected for phospholipids in SLBs, at least on the spatial scales probed (>40nm). Consequently, the large gold nanoparticle tags used in this work do not seem to introduce a change in the overall diffusion mode, but only in the overall mobility. While this kind of effect is expected from theory [21], the magnitude we here report indicates that crosslinking phenomena aside of size are contributing to a great extent. iSCAT experiments on the gold-tagged lipid analogues revealed similar diffusion characteristics, which highlight the viability of the complementary use of both approaches for studying molecular diffusion dynamics, and pave the way for measurements in more complex model and cellular membrane systems. Yet, care has to be taken when using commercially available streptavidin-coated gold nanoparticle samples, since the FCS measurements of the gold nanoparticle tagged lipids disclosed a large amount of streptavidin-only tagged lipids, suggesting the use of dedicated purification protocols.

## MATERIALS AND METHODS

### Lipids

DOPC (1,2-dioleoyl-sn-glycero-3-phosphocholine), DSPE-PEG-Biotin (DSPE: 1,2-distearoyl-sn-glycero-3-phosphoethanolamine, PEG (molecular weight 2000): long linker between lipid and biotin) and DOPE-Cap-Biotin (DOPE: 1,2-dioleoyl-sn-glycero-3-phosphoethanolamine, CAP: short C6-linker between lipid and biotin) were purchased from Avanti Polar Lipids, Atto488-tagged DPPE (1,2-dipalmitoyl-sn-glycero-3-phosphoethanolamine) and DSPE from Atto-Tec, while KK114-tagged DPPE was available as custom-made stock [27]. All lipids were stored at −20°C in chloroform (Sigma) and sonicated briefly prior to their utilization.

### Preparation of Fluorescent Gold Nanoparticles

Streptavidin-coated, 40nm and 20nm diameter gold nanoparticles were purchased from BBI Solutions (Cat. No BA.STP40) as a 3mM solution (OD 10) in 2mM sodium Phosphate, 0.095% NaN_3_ pH7.2 buffer. We tagged these beads with the fluorescent dye Abberior Star 635P, NHS ester (Abberior GmbH) to detect them in FCS as well as iSCAT. We prepared a solution with 3μM gold beads and 10μM NHS ester dye in Phosphate Buffer Saline (PBS buffer). We added a volume equal to 10% of the resulting solution of 1M NaHCO_3_ to trigger the binding reaction between the NHS-ester group of the dye with the amine groups of the streptavidin coating the nanoparticles. The mixture was incubated at room temperature for 1hr. The resulting solution was dialyzed overnight for at least 24hrs in 4l of PBS buffer in cold room at 4°C in a dialysis membrane (Thermo Scientific, 10000 MWCO, cat.no.68100) to eliminate excess dye and possible impurities. We confirmed that the binding of the dye Abberior Star 635P to the streptavidin on the gold nanoparticles did not significantly alter the fluorescence properties of the dye by determining its fluorescence lifetime τ and fluorescence emission maximum *λ_max_* (see Supplementary Material for method description) and triplet state lifetime τ_T_ (Eq. 1), and comparing values on the gold nanoparticles with those determined for free Abberior Star 635-NHS in solution, which were roughly the same (τ = 3.85 ± 0.05ns on gold nanoparticle and 3.76 ± 0.02ns in solution, and *λ_max_* = 655nm and τ_T_ = 5-10μs in both cases).

### Supported Lipid Bilayers

The bilayers were formed by spin-coating a mixture of 1.27mM DOPC dissolved in 1:1 chloroform/methanol solution with the addition of either 0.01mol% KK114-DPPE, or 0.5mol% of either DOPE-Cap-Biotin or DSPE-PEG-Biotin. 25μl of each lipid solution were dropped on acid-cleaned round microscope cover glass (Menzel-Glaser, 25mm diameter, 1.5mm thickness) and positioned on a spin-coater (Chemat Technology). The cover glass was immediately spun at 3000 to 4000 rpm for 30sec. This procedure induced evaporation of the solvent and formation of the SLB on the cover glass, which remained stable for hours. The SLB-coated glass was then placed in a microscopy liquid-tight chamber. Bilayers featuring biotinylated lipids were tagged with the fluorescent gold nanoparticles in this phase, adding a solution of 3μM tagged gold beads and incubating at room temperature for 30min, which was followed by another washing step to remove unbound particles. In a control experiment, we added 10mM of free biotin (Life Technologies, ref. B20656) seconds after the addition of the gold nanoparticles. Samples with KK114-tagged DPPE did not undergo this passage. Atto488-DPPE (Atto-Tec) was used to independently assess the presence of a bilayer region in the sample. The bilayers were kept hydrated during FCS and iSCAT experiments with a 18mM HEPES, 150mM NaCl, pH 7.35 buffer.

### STED-FCS experiments

Confocal and STED-FCS measurement were performed at room temperature on a custom-designed STED microscope, based on an Abberior instrument RESOLFT system (Abberior GmbH) [5], [28], [29]. Fluorescence was excited with a pulsed PicoQuant 640nm diode laser (80 ps pulses, repetition rate 80 MHz, LDH-640), and super-resolution STED microscopy achieved by depleting fluorescence around the center of the excitation spot with 755nm laser light coming from a tunable pulsed source (MaiTai HP titanium-sapphire, repetition rate 80Mhz, Spectra-Newport), whose focal pattern was shaped as a donut with the use of a vortex phase-plate (VPP-1a, RPC Photonics, Rochester, NY). FCS data with an acquisition time of 5sec was recorded using a hardware correlator (Flex02-08D, Correlator.com).

FCS data was obtained through an additional hardware correlator (Flex02-08D, Correlator.com), and analysed using the FoCuS-point fitting software [30]. FCS data for the KK114-DPPE featuring bilayers were analysed using a single component, 2D diffusion model,

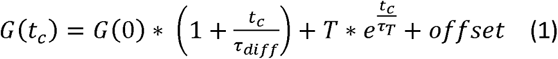

where t_c_ is the correlation time, G(0) and offset are the correlation curve’s amplitude and baseline, respectively, τ_diff_ is the average transit time of the labelled molecules through the observation spot, and T and τ_T_ are amplitude and temporal decay parameters accounting for transient transition of the dyes into the dark triplet state [5], [30]. No anomaly coefficient accounting for possible anomalous diffusion (as has been applied in previous live-cell STED-FCS experiments [12]) was needed to accurately fit the data.

FCS data acquired for the gold nanoparticle tagged lipids were fitted by a two-component model,

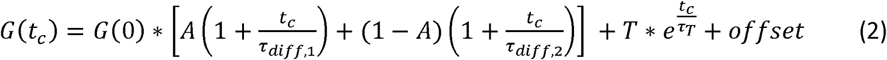

where τ_diff,1_ and τ_diff,2_ are the average transit times of the first and second component, respectively, and A is the relative fraction observed for the first component.

The quality of the fits was evaluated as before [26] by calculating the Akaike information criterion (AIC) for each model according to the formula:

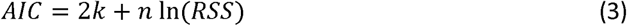

Where k is the number of parameters estimated by the model, n is the number of points and RSS is the residual sum of squares [31]. Note that for the one-component model k = 4, while for the two-component model k = 6. The relative likelihood of these model can be estimated from the AIC by using the relation:

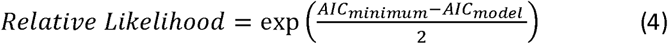

The results of this test indicate the need for using a two- instead of a single-component model in the case of the gold-nanoparticle tagged lipids.

We recorded FCS data for different observation spot diameters d. The full-width-at-half-maximum (FWHM) intensity-based diameter d of the observation spot was reduced from the diffraction-limited, confocal case (d = 240nm, determined from confocal images of 20nm Crimson Beads, Life Technologies) by increasing the power P_STED_ of the STED laser. Measurements on SLBs composed of DOPC with the addition of 0.01mol% KK114-DPPE were performed prior to each experiment session to calibrate the dependence of observation spot size d on the STED laser power, following the relation [5],

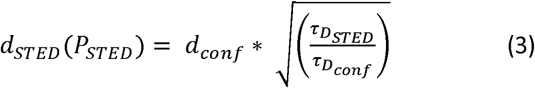

where d_conf_ = 240nm is the FWHM-diameter for the confocal recordings, d_STED_(P_STED_) that for a certain STED power P_STED_, and τ_Dconf_ and τ_DSTED_ the transit times determined for the STED and confocal recordings, respectively. This relation is valid since KK114-DPPE has been shown to diffuse freely in the DOPC-SLBs, i.e. the transit time scales linearly with d^2^ [27].

In the final measurements, we recorded FCS data for three different diameters (confocal and two STED powers) down to 40nm for KK114-DPPE and gold nanoparticle-tagged DOPE-PEG-Biotin and down to 65nm for gold nanoparticle-tagged DOPE-cap-Biotin. We could not record for lower d in the case of DOPE-cap-Biotin due to low signal-to-noise ratios.

Diffusion coefficients D were calculated following the relation:

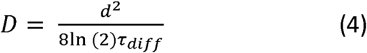

For each sample, we measured correlation curves in at least four different areas of the SLB, repeating the measurements at least five times to ensure repeatability. For each different observation spot, three measurements were carried out: one confocal FCS and two different STED power settings.

### iSCAT experiments

We performed iSCAT experiments on a custom built iSCAT microscope, assembled following the general procedure described in [32]. The output from a 650nm solid-state laser diode (OdicForce lasers) was scanned laterally by two acousto-optic deflectors (AOD, Gooch & Housego and AA Opto-Electronics), sent through a quarter-wave plate and into the back focal plane of a 60x, 1.42NA oil immersion objective (Olympus PlanApo), mounted in inverted geometry. The reflected component from the glass support and the back-scattered light from the sample were collected by the objective, reflected onto the detection path by a polarizing beam splitter, where they interfered, generating the contrast in the final image. Both components were focused on a CMOS camera (Photon Focus MV-D1024-160-CL-8), to acquire images with an effective magnification of 333x (31.8nm pixel size). The imaging plane was stabilized using a Total Internal Reflection fluorescence (TIRF) system described previously [32]. The output from a 462nm solid state laser diode was focused on the back aperture of the objective, and the beam positioned off-centre with a movable mirror until the reflection of the same beam appeared on the other side of the back aperture. This reflection was then deflected through a cylindrical lens (CL) and onto a second CMOS camera (Point Grey Firefly), positioned off-focus with respect to the CL. The position of the resulting line, correlated to the z position of the sample on the microscope stage, was kept constant using a feedback loop that utilized a piezo element (Piezosystem Jena GmbH) to move the objective as required to keep the sample in focus. This system was also exploited to obtain simultaneous TIRF imaging of the lipid bilayers. By adding Atto488-labelled DPPE to the lipid mixture, we could ascertain the presence of the lipid bilayer in the field of view, and that gold beads were located on it.

The sample was illuminated with 2.5mW (60μW/μm^2^) of the 650nm laser light to ensure near-saturation conditions for the scattering signal with an exposure time of 8 ms, and acquired 3 to 5 s long movies with a sampling frequency of 100 Hz. Static features in the movies were eliminated through a temporal median filtering operation, leaving only the moving gold nanoparticles visible ([20], [32], [33]). Single Particle Tracking was performed using the Radial Symmetry algorithm for candidate particle refinement [34] and the u-Track algorithm for the tracking in itself [35], as implemented in the TrackNTrace framework for MatLab [36]. We determined mean-squared-displacement (MSD) plots from each individual single-molecule track and calculated the diffusion coefficient from the ensemble-averaged MSD data using the equation

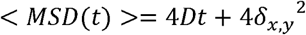

where D is the diffusion coefficient, t is the time difference at which we calculate the MSD, and δ_x,y_ is the localization measurement error that originates from finite localization precision. [37]

## RESULT AND DISCUSSION

We measured the diffusion dynamics of differently tagged fluorescent lipid analogues in a homogeneous (DOPC, 1,2-dioleoyl-sn-glycero-3-phosphocholine) supported lipid bilayer (SLB) (Figure 1a): organic dye (KK114) tagged DPPE (1,2-dipalmitoyl-sn-glycero-3-phosphoethanolamine), and 40-nm gold-nanoparticle tagged DSPE-PEG-Biotin (1,2-distearoyl-sn-glycero-3-phosphoethanolamine, with a long PEG(2000)-linker) and DOPE-Cap-Biotin (1,2-dioleoyl-sn-glycero-3-phosphoethanolamine with a short C6-linker). We employed streptavidin-coated gold nanoparticles which we labelled with an organic dye using an optimized protocol (Materials and Methods). These modified tags target the biotinylated lipids on the SLB surface, and allowed both iSCAT and STED-FCS measurements with high contrast.

### Confocal FCS measurements

We first measured confocal FCS data for the differently tagged fluorescent lipid analogues. Figure 1b shows representative correlation curves, which highlight significant differences between the KK114-and gold nanoparticle tagged lipids. On the one hand, the decay is shifted towards longer correlation times for the gold nanoparticle tagged lipid, revealing decreased mobility. Furthermore, we had to fit the data using different models. While a single diffusing component describes the FCS data of the KK114-tagged lipid well (Eq. 1, with average transit time τ_diff_ = 2.0 ± 0.01 ms, average and standard deviation from 10 measurements, Relative Likelihood of the two component model being favourable is very low 5*10^−172^), we needed to employ a two-component model (Eq. 2) to adequately fit the FCS curves of the fluorescent gold nanoparticle-tagged lipids (Fig. 1b, τ_diff,1_ = 2.5 ± 0.2ms and τ_diff,2_ = 120 ± 30ms for DSPE-PEG-Biotin and τ_diff,1_ = 3.1 ± 0.2ms and τ_diff_,2 = HO ± 30ms for DOPE-Cap-Biotin, average and standard deviation from 10 measurements, Relative Likelihood of the one component model being favourable very low 10^−90^). For fit results see also Supplementary Table 1. The most obvious cause of multiple components in the latter case would be from additional lipids tagged with fluorescent streptavidin only (i.e. lipids with and without a gold bead attached) or additional fluorescent streptavidin-coated gold beads non-specifically bound to the membrane (i.e. not linked to a biotinylated lipid). We ruled out the latter, since we did not observe any fluorescence signal when adding fluorescent streptavidin-coated gold beads to a SLB without biotinylated lipids. On the other hand, we obtained similar average transit times (τ_diff_ = 2.1 ± 0.5ms) for Abberior Star 635-conjugated streptavidin diffusing on a SLB containing biotinylated lipids.

We confirmed that free fluorescent streptavidin was causing the additional diffusing component by studying the composition of the gold nanoparticle solution, by performing Native-PAGE electrophoresis on the gold nanoparticle solution as supplied by the vendor, on our custom fluorescent nanoparticle preparation, and on a sample of pure streptavidin (Supplementary Figure 1 and Supplementary Material). While under our conditions the gold nanoparticles did not diffuse into the gel (producing dark bands at the sample well), but both samples still produced multiple bands (lanes 2 and 3), with the most intense forming in correspondence of the 52 kDa reference band, i.e. the molecular weight of streptavidin. Consequently, more sophisticated purification procedures (e.g. using chromatographic separation approaches) should be used in the future to reduce or hopefully eliminate the contribution of the faster diffusing component from the FCS correlation curve. However, we remark that the streptavidin-only component is only present in FCS measurements (since free streptavidin is fluorescently-tagged by our labelling protocol), and not in the final iSCAT recordings, since the scattering signal from the large gold nanoparticles overwhelms the signal from the smaller streptavidin. Still, even the non-detectable streptavidin might introduce slight bias due to cross-linking of biotinylated lipids.

Because of our controls, the slow component must reveal the mobility of the truly gold nanoparticle tagged lipids. As pointed out, compared to the purely dye-tagged lipid analogue the decays of the FCS data of the gold nanoparticle-tagged lipids are strongly shifted towards longer correlation times highlighting reduced mobility. Analysis of the data (eq. 4) reveals diffusion coefficients of D = 0.12 ± 0.04 μm^2^/s and 0.08 ± 0.02 μm^2^/s for the gold nanoparticle tagged DSPE-PEG-Biotin and DOPE-cap-Biotin lipid analogues, respectively, compared to D = 5.3 ± 0.2 μm^2^/s for the KK114-tagged DPPE lipid analogue.

Consequently, the 40 nm gold nanoparticle tag introduced a roughly 40-to-60-fold slow-down of diffusion. This could result from the increased size introduced by the large tag [19, 22], and/or by cross-linking of several biotinylated lipids through a single bead due to the presence of multiple streptavidin proteins on the gold beads (Figure 1a). We attempted to reduce the latter effect by adding free biotin less than five seconds after the gold nanoparticles after the fluorescent gold nanoparticles) to the solution, in order to block and thus minimize excess binding sites on the nanoparticle surfaces. FCS data taken under this condition still show two diffusing components, where the slower one, belonging to gold nanoparticle-tagged lipids, is compatible to those measured without the introduction of the blocking biotin solution (τ_diff,2_ = 140 ± 40 ms, D = 0.08 ± 0.02 μm^2^/s for DSPE-PEG-Biotin with p=0.2, τ_diff,2_ = 130 ± 40 ms, D = 0.08 ± 0.02 μm^2^/s for DOPE-cap-Biotin with p=0.5). While the presence of free biotin still cannot fully exclude residual cross-linking, this control experiment indicates that the greater size of the gold nanoparticle alone may introduce a slow-down in diffusion, as proposed before [21]. In addition, we tested the effects of the nanoparticle size on the general mobility. For this, we carried out FCS experiments on the same SLB sample using fluorescent, streptavidin-coated 20nm gold nanoparticles. Compared to the 40nm gold nanoparticles, we expect the decrease in size to result in a reduced number of potential cross-linking sites and overall in a less pronounced slow-down in diffusion. Indeed, the diffusion coefficients measured in this instance revealed a slightly increased mobility (D = 0.3±0.1 μm^2^/s for DSPE-PEG-Biotin and D = 0.2±0.1 μm^2^/s for DOPE-cap-Biotin, i.e. only 20-25-fold reduction compared to KK114-tagged DPPE) relative to the 40 nm large gold nanoparticle tagged lipids (D = 0.12 ± 0.04 μm^2^/s and 0.08 ± 0.02 μm^2^/s, respectively, i.e. 40-to-60-fold reduction compared to KK114-tagged DPPE), further highlighting the fundamental influence of the tag’s characteristics on the diffusion of the biotinylated lipids.

Comparing the diffusion characteristics of the gold nanoparticle tagged DSPE-PEG-Biotin and DOPE-cap-Biotin lipids, it seems that the longer PEG linker (compared to the C6-linker of the cap-biotin) slightly reduces the biasing effect of the gold nanoparticle tag. We can assume that the different saturation degrees of the DSPE and DOPE lipids did not introduce the 1.5-fold difference in mobility, since no significant difference in their overall mobility was previously reported on model membranes [5] as well as from our own FCS measurements of Atto488 dye -conjugated DSPE and DOPE lipids on the same SLB system, which revealed similar values for both (D = 4.6±0.3 μm^2^/s for ATTO488-DSPE and D = 4.3±0.2 μm^2^/s for Atto488-DOPE).

### STED-FCS measurements

We also aimed to investigate whether tagging with a large gold nanoparticle only changes the overall mobility or also the lipid’s diffusion mode, i.e. whether it introduced anomalous diffusion. We therefore recorded FCS data for different observation spot diameters (d) as generated by increasing STED laser powers in the STED microscope (STED-FCS, Figure 1c). We expect that, for a freely diffusing phospholipid on a homogeneous DOPC SLB, the value of the apparent diffusion coefficient D should be independent of the observation spot size d. Any variation indicates hindered diffusion, as highlighted as trapped or compartmentalized/hop diffusion following a decrease or increase of D-values towards smaller diameters d, respectively [27]. We indeed observed an independence of D on the observation spot diameter d for the KK-114-tagged DPPE analogue (Figure 1d). More importantly, we also observed a roughly constant D(d) dependency, i.e. purely free diffusion for the gold nanoparticle tagged lipids (at least in the spatial scales tested). Consequently, the gold nanoparticle tags introduced an overall slow-down in mobility, which still remains a free-diffusion character, i.e. no trapping interactions or confinements.

The latter hypothesis was strengthened by recording STED-FCS data of 40 nm gold nanoparticles on a less fluid SLB, composed at 90% of DOPC and 10% Cholesterol. No observable anomaly was observed in this system either, suggesting that the main factor in the reduced diffusivity of lipids are, indeed, the characteristic of the nanoparticle tags. However, the magnitude of the diffusion coefficients for gold tagged lipids (D_conf_ = 0.14±0.07 μm^2^/s, D_STED_ = 0.14±0.04 *μm^2^/s* for DSPE-PEG-Biotin; D_conf_ = 0.12±0.08 μm^2^/sec, D_STED_ = 0.11±0.08 μm^2^/s for DOPE-cap-Biotin) was in this case reduced by 30-to-40-fold reduction in mobility compared to a fluorescent lipid analogue diffusing on a SLB with the same lipid composition (D_conf_ = 4.0±0.2 μm^2^/sec, D_STED_ = 4.0±0.2 μm^2^/s for KK114-DPPE).

A possible artefact influencing the FCS measurements at reduced observation spots, i.e. under STED laser conditions, could be altered photophysical characteristics of the fluorescent dye Abberior Star 635, like fluorescence lifetime or emission spectrum, due to proximity to the gold surface. Such difference would (as shown in [38]) in turn lead to different confinement characteristics of the effective fluorescence observation spot for the measurements on the fluorescent gold-tagged lipids, compared to the dye-only conditions. Therefore, the assumed observation spot diameters d and thus the D(d) dependency would not be comparable across the different experimental conditions. However, we did not observe significant changes in both fluorescence spectrum and lifetime between sole Abberior Star 635 and the Abberior Star 635-streptavidin-gold nanoparticle construct; the fluorescence emission of both was characterized by a peak at 655 nm and lifetimes of 3.76 ± 0.02ns for the free dye and 3.85 ± 0.05ns for the gold nanoparticle–dye conjugate (Materials and Methods and Supplementary Material), respectively. Another source of bias could be optical trapping of the beads at the increased STED laser powers, leading apparently enhanced average transit times and reduced diffusion coefficients. However, no such effect has been observed.

### Interferometric scattering-based Single Particle Tracking

Finally, we complemented our STED-FCS measurements using iSCAT microscopy to track the gold nanoparticle tagged lipids diffusing on the same SLBs and under the same conditions as for the STED-FCS recordings. The gold-tagged lipids appear as dark spots on a white background, due to the nature of image formation in iSCAT (Supplementary Figure 2). No such structures were detected in non-labelled or KK114-DPPE labelled SLBs, due to the lack of moving objects with sufficiently large scattering cross-section. The maximum spatial localization precision of the set-up was determined to be 2.6 nm, measured on 40nm gold nanoparticles. This value was determined by measuring the FWHM of the distribution of the relative positions of two non-moving particles their relative positions. The temporal resolution of these measurements, given by the inverse of the sampling rate, was 0.01 seconds. Consequently, the movies allowed the recording of molecular tracks of individual lipid analogues with high spatial and temporal resolution. We analysed 97 and 63 tracks for the DOPE-cap-Biotin and DSPE-PEG-Biotin samples, respectively, and calculated the ensemble average mean-squared-displacement (MSD) plots (MSD versus lag time τ) (Figure 2a,b). The single tracks were not considered in their individuality, as we were at this stage interested only in the possibility of comparing this method with (STED-)FCS, a method that inherently averages all multiple particle transits. The ensemble-averaged MSD plots for both gold nanoparticle tagged lipids were well described as free diffusion [37] with diffusion coefficients D = 0.31 ± 0.01 μm^2^/s for DSPE-PEG-Biotin and D = 0.17 ± 0.005 μm^2^/s (DOPE-cap-Biotin). This was confirmed when plotting values of MSD/4τ versus lag time τ, which shows a constant dependence as expected for free diffusion (Inserts in Fig.2a,b). The results from the iSCAT recording were in rough agreement with the slower components from the STED-FCS recordings (D = 0.12 ± 0.04 μm^2^/s for DSPE-PEG-Biotin and D = 0.08 ± 0.02 μm^2^/s for DOPE-cap-Biotin). The differences in the absolute values of the apparent diffusion coefficients as determined by the two methods may be attributed to the differences in the experimental methods employed [26], [39]. Nevertheless, these values are in the same order of magnitude, and the iSCAT recordings reveal a similar difference in mobility between the two biotinylated lipid species as for STED-FCS.

**Figure 2.**
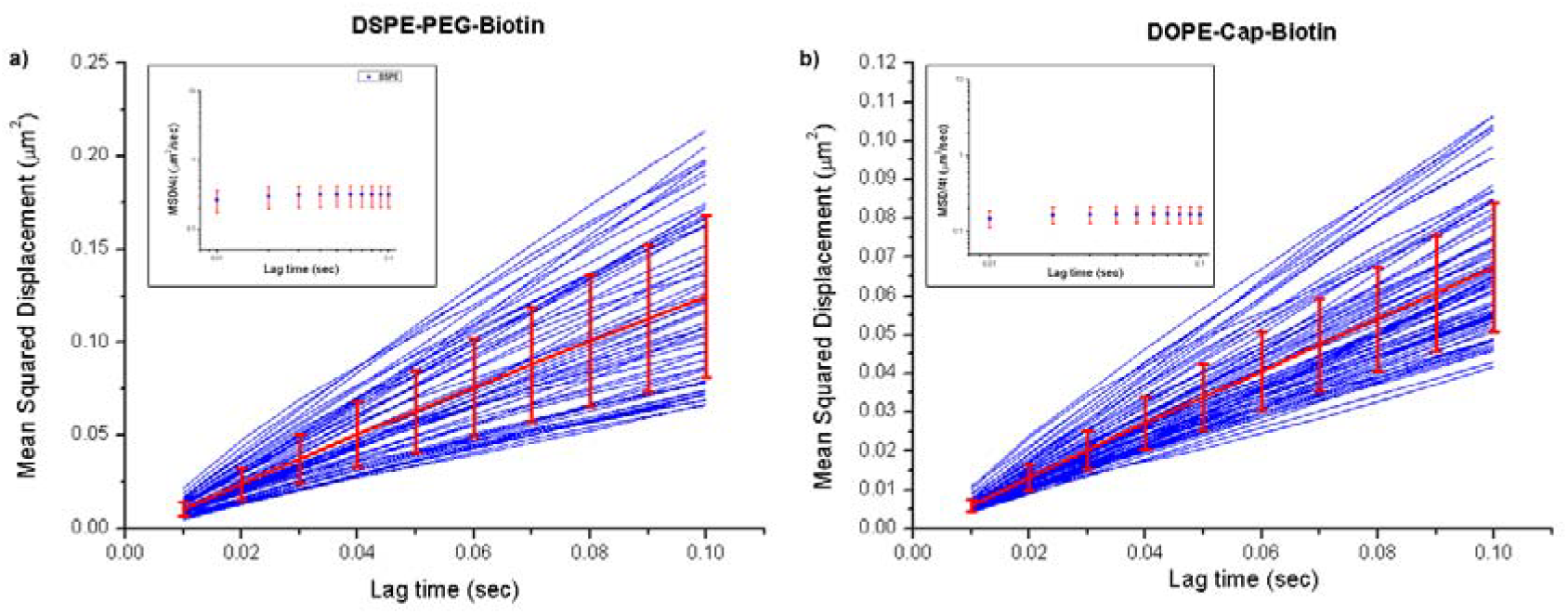
Representative iSCAT microscopy data of the diffusion of the gold nanoparticle tagged (a) DSPE-PEG-biotin and (b) DOPE-cap-biotin lipids on the SLB: Mean-Squared-Displacement (MSD vs trajectory lag time t) plots calculated for 63 and 97 different respective trajectories (blue: individual MSD data, red: ensemble average with error bars from standard-deviations-of-the-mean) and linear fits against average MSD = 4Dt (red) with resulting values of D. (Inserts) Plots of the ensemble average MSD/4t over t for both diffusing species, highlighting a free diffusion behaviour.

## Conclusions

The aim of this work was to compare the performance of two experimental techniques, STED-FCS and iSCAT-based SPT, and explore the viability of protein-coated gold nanoparticles as tags for SPT by detecting the diffusion characteristics of gold nanoparticle-tagged biotinylated lipids on simple, homogeneous model membranes. This simple case study was designed such that we could readily delineate the effect on the mode of diffusion and on the magnitude of the diffusion coefficients for gold-particle tagged lipids (as compared to dye tagged lipids) in SLBs.

Our complementary STED-FCS and iSCAT measurements reveal several important issues. First and foremost, we have observed that the streptavidin-coated 40nm gold nanoparticle tag introduced a 40-to-60-fold slow-down in diffusion. The source of this effect could potentially be either the sheer size of the gold nanoparticle, probe induced cross-linking, or both. However, while existing studies acknowledge that the dimensions of such a probe have a significant effect on the diffusion rate of the tagged particles, through steric hindrances, these effects are usually expected to be much less than those reported in this work (less than 10-fold) but difficult to quantify absolutely [21,41]–[43]. We therefore conclude that a large influence on the general mobility results from the cross-linking of several biotinylated lipids by the gold nanoparticle tags. We attempted to dampen this effect with the addition of free biotin, to saturate the excess binding sites of the gold nanoparticle, but this proved ineffective. It is therefore evident, as proposed in [21], that an additional purification step has to be undertaken, to ensure that the majority of the employed probes are monovalent. Interestingly, we have also noticed that using smaller gold nanoparticles (20nm) produced roughly a 2 to 3-fold increase in the observed diffusion coefficient compared to the 40nm large tags. This may, besides the reduction in sheer size, be due to a reduced potential for cross-linking related to a smaller available linkage area. Nevertheless, lipids tagged with gold nanoparticles still showed free diffusion (at least on the spatial scales probed > 40 nm) making these nanoparticle probes still potentially usable for studies of membrane diffusion modes.

Furthermore, we have shown that great care has to be taken when using commercially available streptavidin-coated gold nanoparticle samples, since FCS measurements and Native PAGE electrophoresis of the gold nanoparticle tagged lipids disclosed a large amount of streptavidin-only tagged lipids, which produce a much smaller signal in the iSCAT measurements, but might still introduce other bias such as additional cross-linking. Finally, while iSCAT-SPT measurements revealed larger absolute values of diffusion coefficients compared to STED-FCS, both approaches picked up consistent relative differences in mobility between lipid analogues with different tag-linkers (cap vs. PEG) and saturation degrees of the lipid chains (DSPE vs. DOPE).

Our results highlight the validity of the complementary use of both approaches (STED-FCS and iSCAT) for studying molecular diffusion dynamics. Yet, while gold-nanoparticle tagged lipids will most probably accurately report on the diffusion mode, they will do so with a much lower absolute diffusion coefficient compared to organic-dye tagged lipids. The ultimate test will be the comparison of the different analysis and tagging approaches in more complex cellular membrane systems. Previous experiments on live-cell diffusion dynamics of dye-tagged lipid analogues have given complementary results for STED-FCS and SPT [43]. Further optimization will involve the use of smaller gold nanoparticles, monovalent linkers between the gold tag and lipid, accurate purification protocols, and the application of these same protocols for live-cell measurements. It will then be possible to access the complementary information from STED-FCS and iSCAT-based SPT data to disclose novel details of molecular membrane dynamics.

## Acknowledgements

We thank Dominic Waithe for support on the data analysis on the sample preparations. F. R. and C. E. greatly acknowledge the EPSRC for supporting the DPhil project of F. R. within the Oxford-Nottingham BioImaging Centre of Doctoral Training (ONBI-CDT). Further, we acknowledge support by the EPA Cephalosporin Fund (Bio-iSCAT project), the MRC (grant number MC_UU_12010/unit programs G0902418 and MC_UU_12025), the Wellcome Trust (grant 104924/14/Z/14 and Strategic Award 091911 (Micron)), MRC/BBSRC/EPSRC (grant MR/K01577X/1), Deutsche Forschungsgemeinschaft (Research unit 1905 “Structure and function of the peroxisomal translocon”), the Wolfson Foundation (for initial funding of the Wolfson Imaging Centre Oxford), and the John Fell Fund.

